# Bacterioplankton Diversity and Distribution in Relation to Phytoplankton Community Structure in the Ross Sea surface waters

**DOI:** 10.1101/2021.06.08.447544

**Authors:** Angelina Cordone, Giuseppe D’Errico, Maria Magliulo, Francesco Bolinesi, Matteo Selci, Marco Basili, Rocco de Marco, Maria Saggiomo, Paola Rivaro, Donato Giovannelli, Olga Mangoni

## Abstract

Primary productivity in the Ross Sea region is characterized by intense phytoplankton blooms whose temporal and spatial distribution are driven by changes in environmental conditions as well as interactions with the bacterioplankton community. Exchange of exudates, metabolism by-products and cofactors between the phytoplankton and the bacterioplankton communities drive a series of complex interactions affecting the micronutrient availability and co-limitation, as well as nutrient uptakes in Antarctic waters. Yet, the number of studies reporting the simultaneous diversity of the phytoplankton and bacterioplankton in Antarctic waters are limited. Here we report data on the bacterial diversity in relation to phytoplankton community in the surface waters of the Ross Sea during the austral summer 2017. Our results show partially overlapping bacterioplankton communities between the stations located in the Terra Nova Bay coastal waters and the Ross Sea open waters, suggesting that the two communities are subjected to different drivers. We show that the rate of diversity change between the two locations is influenced by both abiotic (salinity and the nitrogen to phosphorus ratio) and biotic (phytoplankton community structure) factors. Our data provides new insight into the coexistence of the bacterioplankton and phytoplankton in Antarctic waters.

## 1 Introduction

Antarctica and the Southern Ocean are central to Earth’s climate and oceanic circulation systems (Convey and Peck, 2019). Primary productivity in this region is characterized by intense phytoplankton blooms whose temporal and spatial distribution are driven by different environmental conditions, although the mechanisms regulating these processes are still poorly known (Moore and Abbott, 2000; Deppeler and Davidson, 2017). In this context, the Ross Sea is one of the most productive sectors in Antarctica (Smith and Gordon, 1997; Arrigo et al., 1999), with an annual productivity averaging ∼180 g C m^-2^ yr^-1^ (Arrigo et al., 2008). Phytoplankton blooms here have been well studied, with the dominance of diatoms and haptophytes presenting different temporal and spatial patterns (Smith et al., 2014; Mangoni et al., 2017). In the last years, however, changes in phytoplankton blooms and dynamics that contrast the classical Antarctic paradigm have been observed (DiTullio et al., 2000; Phan-Tan et al., 2018; Mangoni et al., 2019; Bolinesi et al., 2020b), suggesting that phytoplankton-bacteria interactions play a key role in structuring the trophodynamics in this area (Bertrand et al., 2007, 2015).

The assumption that the Antarctic Ocean microbial communities were generally species poor, has been re-discussed in recent years, since many results suggested that microbial diversity is significantly higher than previously recognised (Murray and Grzymski, 2007; Wilkins et al., 2013; Silvi et al., 2016). In the last decade, owing to technological and analytical improvement, researched have demonstrated that in pelagic marine food webs, prokaryotes significantly contribute to the structuring of the micro-eukaryotic community (Kirchman et al., 2001; Wilkins et al., 2013; Delmont et al., 2014). Prokaryotic communities contribute to or dominate several key ecosystem processes, including primary production, the turnover of biogenic elements, the mineralization of the organic matter and the degradation of xenobiotics and pollutants (Cole, 1982; Azam et al., 1983; Croft et al., 2005; Azam and Malfatti, 2007; Falkowski et al., 2008; Sher et al., 2011).

Early work in Antarctic waters revealed that bacterial biomass might represent up to 30 % of total microbial biomass in coastal areas (Fiala and Delille, 1992). In Antarctica, free-living marine microbial community composition can differ significantly between locations at relatively small spatial and temporal scales, responding to environmental variations of temperature, salinity, nutrients, or the presence of oceanic fronts (Lozupone and Knight, 2007; Nemergut et al., 2011; Raes et al., 2018). For example, in Terra Nova Bay a substantial difference in terms of bacterial assemblages has been observed between coastal and offshore stations and along the water column (Celussi et al., 2009). Heterotrophic members of the *Alphaproteobacteria* and *Gammaproteobacteria* class of the *Proteobacteria* are reported as dominant phylotypes in Antarctic waters (Giovannoni et al., 2005; Murray and Grzymski, 2007; Wilkins et al., 2013), and studies in this and other marine ecosystems indicate that bacterial growth is frequently dependent on phytoplankton-derived DOM (Church et al., 2000; Morán et al., 2001, 2002; Piquet et al., 2011; Ducklow et al., 2012; Kim et al., 2014). A broad diversity amongst class *Flavobacteria* has been also reported in different sub-areas of the Southern Ocean (Abell and Bowman, 2005). According to Piquet et al., (2011) melt water stratification and the transition to non stabilized Antarctic surface waters may have an impact not only on micro-eukaryotes but also on bacterial community composition, with a shift from an *Alpha-* and *Gammaproteobacteria* to a *Cytophaga–Flavobacterium–Bacteroides*-dominated community under mixed conditions. Some studies found an unusual presence of strictly anaerobic *Epsilonproteobacteria* (now reclassified as phylum *Campylobacterota*) in the bottom of sea ice (Gentile et al., 2006), probably as a consequence of the oxygen decay and sulfide accumulation, caused by high degradation rates of sympagic diatoms by aerobic and anaerobic heterotrophs (Brierley and Thomas, 2002).

Shifts in the eukaryotic community have been often reported as rapidly followed by a shift in the bacterial community (Billen and Becquevort, 1991; Piquet et al., 2011) with a series of phytoplankton–bacterial interactions resulting in both positive and negative feedback loops (Bertrand et al., 2015). For example, in temperate waters phytoplankton blooms in spring and summer induce changes in bacterioplankton community structure (Fuhrman et al., 2006; Teeling et al., 2016; Chafee et al., 2018). These community dynamics have been shown to be recurrent, indicating a phytodetritus-driven seasonality and suggesting that the phytoplankton-prokaryotic interactions in surface waters are more sophisticated than previously thought (Seymour et al., 2017). Mechanisms addressing the nature of the mutualistic interaction between phytoplankton and bacterioplankton communities have been proposed over time. For example, the exchange of phytoplankton exudates and bacteria-produced cobalt containing vitamin B_12_ represents one of the best studied feedback loops (Bertrand et al., 2007, 2011; Bolinesi et al., 2020b). The production and release of vitamin B_12_ by bacteria, in fact, depends on the degradation of phytoplankton exudates, establishing a complex feedback mechanism between prokaryotic and phytoplanktonic communities by heterotrophic bacteria (Fang et al., 2017). Similar trophic interaction between phytoplankton and bacterioplankton in the marine ecosystem, and especially in polar regions, might be more common than previously known. The bacterial-phytoplankton interaction may thus evolve in a series of complex relationship, affecting directly or indirectly the micronutrient availability and co-limitation (e.g. iron, cobalt, vitamin B_12_) (Bertrand et al., 2007; Tagliabue et al., 2017; Bolinesi et al., 2020a) as well as nutrient uptake (Amin et al., 2015; Bertrand et al., 2015). Yet, despite this the number of studies reporting the simultaneous diversity of the phytoplankton and bacterioplankton community in the Antarctic waters are comparatively few (Di Poi et al., 2013; Flaviani, 2017; Richert et al., 2019). In this study we analyzed the bacterial diversity in relation to phytoplankton in the sub-surface waters of Terra Nova Bay and Ross Sea, providing new insight into the coexistence of the two communities in Antarctic waters.

## 2 Material and Methods

### Sampling procedure and study site

Seawater samples were collected at different depths in correspondence of deep chlorophyll maximum, or at ∼20 m depth where the fluorescence profile showed a rather homogeneous distribution of chlorophyll-a in the upper layer (Table 1). Sampling activities were carried out on the R/V Italica during the Austral Summer 2017, in the framework of P-ROSE (Plankton biodiversity and functioning of the Ross Sea ecosystems in a changing Southern Ocean) and CELEBeR (CDW Effects on glacial mElting and on Bulk of Fe in the Western Ross sea) projects - Italian National Antarctic Program - funded by the Ministry of Education, University and Research (MIUR). The area of investigation falls within two different zones (Figure 1) of the Ross Sea, the coastal area of Terra Nova Bay (TNB) and the Ross Sea Open Water (RSOW). Water samples were collected using a carousel sampler (Sea-Bird Electronics 32) equipped with 24 12-L Niskin bottles and a conductivity–temperature-depth (CTD) instrument (9/11 Plus; Sea-Bird Electronics), along a transect from the coastal area of Terra Nova Bay to the oper Ross Sea (Figure 1C), crossed by a north-south aligned secondary transect carried out in the RSOW area. For the bacterial diversity analysis, at each station 500 mL of seawater were collected from the Niskin bottle and filtered onto a 0.22 μm filter (Whatman, 47 mm diameter) successively stored at −80 °C and transported back to the lab. For the analysis of total phytoplankton biomass, 500 mL of sea water were filtered on board through 0.45 µm GF/F filter (Whatman, 47 mm diameter) and preserved frozen at −80°C. For the determination of phytoplankton functional groups by chemotaxonomic criteria, 2 L of seawater were filtered on board onto 0.45 µm GF/F filter (Whatman, 47 mm diameter) and stored at −80 °C. For the phytoplankton counts and taxonomic analysis, water samples were immediately fixed with 4 % CaCO_3_ buffered formalin solution. For the analyses of macronutrient concentrations (NO_3_ ^-^, NO_2_ ^-^, NH_4_ ^+^, Si(OH)_4_, PO_4_ ^3-^), water samples were taken directly from the Niskin bottles and stored at −20 °C in 20 mL low-density polyethylene containers until laboratory analysis.

**Figure 1.**
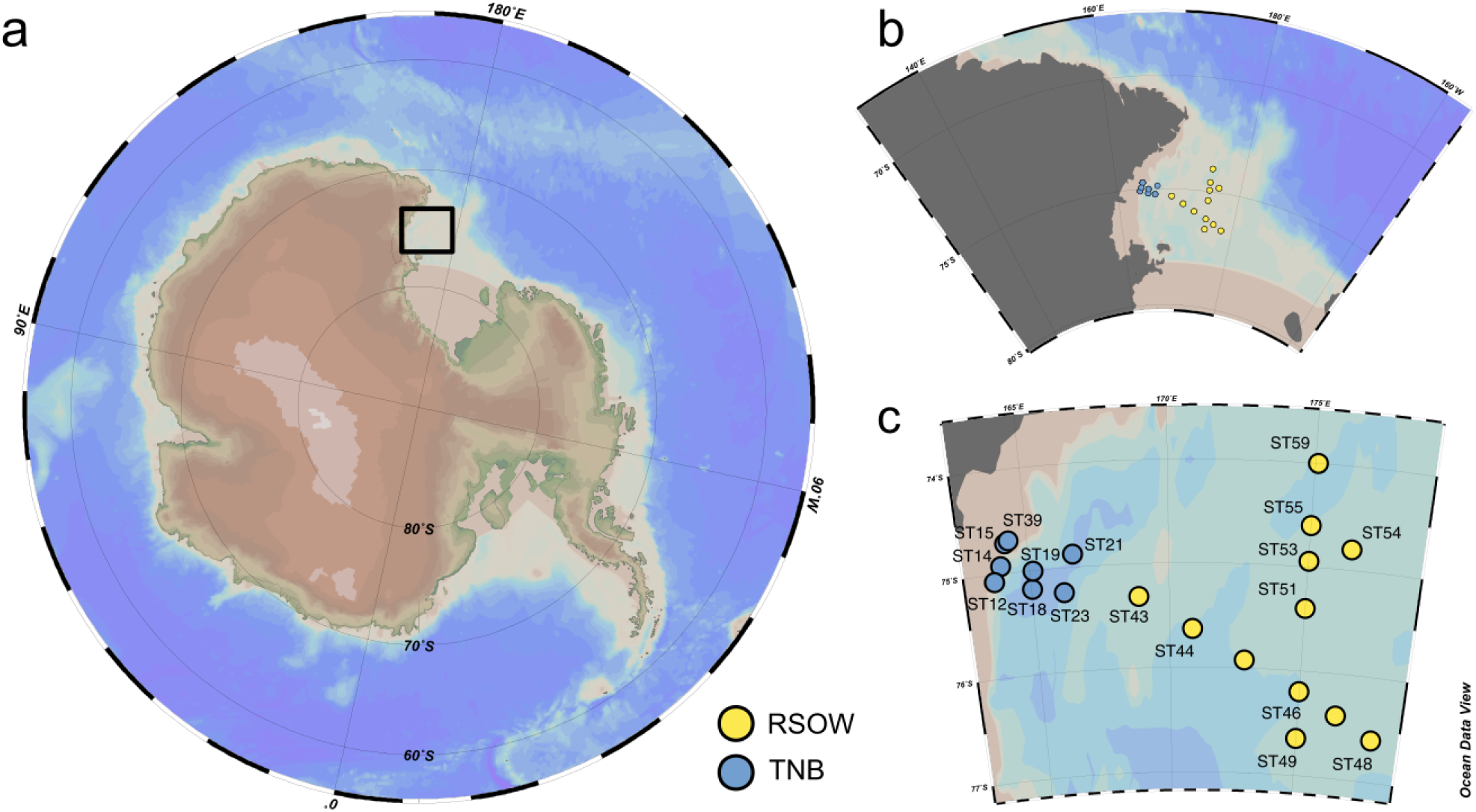
Study location and distribution of the sampled stations across the Ross Sea. a) Location of the study site within the ross sea. b and c) details of the spatial distribution of the sampled stations and their division in Terra Nova Bay (TNB) stations proximal to the sea-ice border and the coast and the Ross Sea Open Waters (RSOW) stations, located further off-shore. This division and color scheme is consistent throughout the paper.

**Table 1.** Summary of the sampled stations and main environmental variables.

| Station | Sampling area | Latitude (°N) | Longitude (°E) | Bottom depth (m) | Sampling depth (m) | Salinity (PSU) | Temperature (°C) | Chlorophyll-a (µg/L) |
| --- | --- | --- | --- | --- | --- | --- | --- | --- |
| ST12 | TNB | -75.07186 | 163.70473 | 867 | 35 | 34.46 | -0.9 | 2.5 |
| ST14 | TNB | -74.9274 | 163.9883333 | 342 | 20 | 34.28 | 1.7 | 3.8 |
| ST15 | TNB | -74.71124 | 164.23106 | 498 | 20 | 34.36 | 1.7 | 2.4 |
| ST18 | TNB | -75.17703 | 165.04118 | 1,054 | 15 | 34.1 | 1 | 2.4 |
| ST19 | TNB | -75.0045667 | 165.1273167 | 925 | 20 | 34.67 | 0.3 | 2.4 |
| ST21 | TNB | -74.8748767 | 166.5811933 | 886 | 30 | 34.1 | -0.1 | 1.3 |
| ST23 | TNB | -75.23705 | 166.1794517 | 852 | 15 | 34 | 0.5 | 2 |
| ST39 | TNB | -74.7128833 | 164.2245833 | 497 | 25 | 34.48 | -0.4 | 1.3 |
| ST43 | RSOW | -75.3133617 | 168.8873467 | 351 | 28 | 34.18 | -0.2 | 1.7 |
| ST44 | RSOW | -75.6297333 | 170.8507833 | 568 | 39 | 34.42 | -0.6 | 1.1 |
| ST45 | RSOW | -75.9229083 | 172.8276267 | 571 | 28 | 34.37 | -0.1 | 0.7 |
| ST46 | RSOW | -76.1994917 | 174.9960983 | 568 | 28 | 34.42 | -0.4 | 1.6 |
| ST47 | RSOW | -76.40035 | 176.4959233 | 409 | 45 | 34.45 | -0.4 | 1.2 |
| ST48 | RSOW | -76.6003167 | 177.9980833 | 292 | 42 | 34.47 | -0.6 | 1.4 |
| ST49 | RSOW | -76.6460167 | 174.9964333 | 425 | 26 | 34.33 | -0.2 | 2.4 |
| ST51 | RSOW | -75.4 | 175.0048333 | 297 | 33 | 34.32 | 0 | 0.3 |
| ST53 | RSOW | -74.9452 | 175.0106 | 326 | 35 | 34.17 | -0.2 | 1.1 |
| ST54 | RSOW | -74.8008333 | 176.5028 | 299 | 28 | 34.28 | -0.1 | 1.7 |
| ST55 | RSOW | -74.6025833 | 175.0045167 | 439 | 15 | 34.15 | -0.3 | 2.2 |
| ST59 | RSOW | -73.99957 | 175.10031 | 579 | 34 | 34.17 | -0.4 | 2 |

### Phytoplankton community structure and inorganic nutrient concentration

Frozen samples were processed in Italy for the determination of chlorophyll-a (Chl-a) and phaeopigments (Phaeo-a) content using a solution of 90 % acetone according to Holm-Hansen, (1956), with a spectrofluorometer (Shimadzu) checked daily with a Chl-a standard solution (from Sigma-Aldrich). HPLC pigments separation was made on an Agilent 1100 HPLC according to the method outlined in Vidussi et al. (1996) as modified by Mangoni et al. (Mangoni et al., 2017). The phytoplankton species composition and cells abundance were determined following the Uthermöhl method (Utermöhl, 1931), according to which at least 400 cells were counted per sample with an inverted light microscope (LM) Zeiss Axiovert Observerz. 1 at 400× magnification. The analyses of inorganic nutrients were performed using a five-channel continuous flow autoanalyzer (Technicon Autoanalyser II), according to the method described by Hansen et al., (1983) which was adapted to the present instrumentation.

### Community DNA extraction

Total community DNA was extracted as previously reported (Giovannelli et al., 2013) with slight modifications. Briefly, each filter was washed with 1.7 ml of extraction buffer solution (100 mM NaCl (pH 8.0), 20 mM EDTA, 50 mM Tris-HCl (pH 8.0)) added with 10 μl Proteinase K (1 mg/ml) and incubated at 37 °C for 1 hour with occasionally mixing. A 100 μl volume of 10 % sodium dodecyl sulfate (SDS) was added to each sample, followed by incubation at 55 °C for 1 hour. The liquid phase was collected and centrifuged for 15 minutes at 10,000×g to separate cellular debris from nucleic acids. The supernatant was extracted twice with an equal volume of phenol, followed by a precipitation with 2.5 volume of ethanol 100% and 0.1 volumes of sodium acetate (3 M) for 12 hours at −20 °C. The DNA was collected by centrifugation, washed in 70 % cold ethanol, air dried, and resuspended in 50 μl of sterile distilled water. The integrity of DNA was assessed spectrophotometrically and by PCR amplification of the 16S rRNA gene using the primes Ribo-For (5’-AGTTTGATCCTGGCTCAG-3’) and Ribo-Rev (5’-CCTACGTATTACCGCGGC-3’) (Fakhry et al., 2008). The PCR products were visualized on 1 % agarose gel stained with ethidium bromide.

### 16S rRNA gene sequencing

Partial 16S rRNA gene sequences were obtained using primer pair Probio_Uni / Probio_Rev, which target the V3 region of the 16S rRNA gene sequence (Milani et al., 2013). 16S rRNA gene sequencing was performed using a MiSeq platform (Illumina) at the DNA sequencing facility of GenProbio srl (www.genprobio.com) according to the protocol previously reported (Milani et al., 2013).

### Bioinformatics and statistical analyses

All sequences were imported in R, and analyzed with the DADA2 package (Callahan et al., 2016). Following the package guidelines, quality plots were performed to check the sequences’ quality. Post-QC reads were trimmed using the filterAndTrim command (truncLen=c(155,145), maxN=0, maxEE=c(2,2), truncQ=2, rm.phix=TRUE, trimLeft=17, trimRight=15). After this step, a parametric error model, based on the convergence between the estimation of error rate and the inference of the sample composition, was performed. Paired-end reads were merged and exact Amplicon Sequence Variants (ASVs) inferred using the dada algorithm. Chimeric sequences were removed and prokaryotic taxonomy assigned using the naive Bayesian classifier method against the Silva Database (v132; https://www.arb-silva.de/documentation/release-132/). ASVs abundance table obtained from DADA2 was further processed in R using *Phyloseq, Vegan,* and *Microbiome* packages (McMurdie and Holmes, 2013; Oksanen et al., 2020; Lahti and Shetty, 2017). Sequences are available through the European Nucleotide Archive (ENA) with bioproject accession number ERP129169. A complete R script containing all the steps to reproduce our analysis is available at https://github.com/giovannellilab/Cordone_et_al_Ross_Sea with DOI: https://doi.org/10.5281/zenodo.4784454.

After the phyloseq object creation, low abundance ASVs (less than 3 reads across the dataset), Mitochondria, Chloroplast, Eukaryotes sequences, and potential contaminants (Sheik et al., 2018) were removed. The resulting dataset was represented by 703 unique ASVs and 535,009 reads. ASVs counts were normalized to the median library size across the dataset. Diversity analyses were carried out using the phyloseq package (McMurdie and Holmes, 2013). The alpha diversity was investigated using both the Simpson and Shannon diversity index among the two sampled areas. The beta diversity was investigated using the UNIFRAC and Jaccard diversity index as implemented in the vegan package (Oksanen et al., 2012). Botht the abundance weighted and unweighted version of the index were used. The resulting similarity matrix was plotted using non-metric multidimensional scaling techniques implemented in the ordination command of phyloseq. The resulting ordination was used to investigate correlations with environmental and phytoplankton variables using the *envfit* and *ordisurf* functions in vegan. Collinearity among the predictors was checked using a Pearson correlation matrix. The linearity of the correlation between the rate of change in the beta diversity and the variables identified as significant by the *envfit* function was checked by plotting the nMDS axis against the variable. Statistically significant differences in the distribution of abundant bacterial genera was tested using the Chi-square test. Co-correlation networks were calculated as a pairwise distribution of each ASV across the entire dataset using Spearman rank correlation and different ρ cutoff selected for network plotting using the igraph package (Csardi and Nepusz, 2006).

## 3 Results

### Phytoplankton community and physical-chemical properties of the water column

Water temperature ranged between −0.3 °C and 1.7 °C (mean value of 0.7±0.8 °C) in TNB, and between −0.6 °C and 0.1 °C (mean value of −0.3±0.2 °C) in the RSOW. (Kruskal-Wallis, p<0.05). Salinity ranged between 33.98 and 34.39 (mean of 34.2±0.2) in TNB, while showed less variability in RSOW with values ranging between 34.2 and 34.4 (mean of 34.3±0.1) (Kruskal-Wallis, non significant). Further details on physical water column properties are described in T/S diagrams (Bolinesi et al., 2020b) highlighting differences between sud-systems. In TNB, dissolved inorganic nitrogen (DIN) concentrations, as sum of nitrate, nitrite and ammonium, ranged between 11.09 µmol/L and 26.37 µmol/L, PO_4_^3-^ ranged between 0.79 µmol/L and 1.82 µmol/L, while Si(OH)_4_ ranged between 35.77 and 49.95 µmol/L. In RSOW DIN ranged between 18.61 µmol/L and 25.52 µmol/L, PO_4_^3-^ between 1.24 and 1.63 µmol/L and Si(OH)_4_ between 42.41 µmol/L and 55.68 µmol/L (Table 2).

**Table 2.** Concentrations of nutrients in the sampled stations.

| Station | Sampling area | DIN<br>($\mu\text{mol/L}$) | $\text{PO}_4^{3-}$<br>($\mu\text{mol/L}$) | $\text{Si(OH)}_4$ ($\mu\text{mol/L}$) | N/P |
| --- | --- | --- | --- | --- | --- |
| ST12 | TNB | 26.37 | 1.82 | 39.78 | 14.49 |
| ST14 | TNB | 11.09 | 0.83 | 35.77 | 13.37 |
| ST15 | TNB | 11.81 | 0.81 | 37.62 | 14.6 |
| ST18 | TNB | 12.32 | 0.79 | 37.8 | 15.68 |
| ST19 | TNB | 17.47 | 1.23 | 49.48 | 14.22 |
| ST21 | TNB | 21.07 | 1.46 | 49.95 | 14.44 |
| ST23 | TNB | 15.16 | 0.89 | 39.99 | 17.05 |
| ST39 | TNB | 22.43 | 1.63 | 45.52 | 11.64 |
| ST43 | RSOW | 22.01 | 1.29 | 42.41 | 16.45 |
| ST44 | RSOW | 23.35 | 1.57 | 48.88 | 14.01 |
| ST45 | RSOW | 22.18 | 1.63 | 51.41 | 13.21 |
| ST46 | RSOW | 18.61 | 1.62 | 55.68 | 10.65 |
| ST47 | RSOW | 16.79 | 1.47 | 50.3 | 10.37 |
| ST48 | RSOW | 18.26 | 1.49 | 49.97 | 11.1 |
| ST49 | RSOW | 19.67 | 1.42 | 53.77 | 13.25 |
| ST51 | RSOW | 20.58 | 1.59 | 54.87 | 12.2 |
| ST53 | RSOW | 24.23 | 1.24 | 47.08 | 18.65 |
| ST54 | RSOW | 23.83 | 1.48 | 45.41 | 14.47 |
| ST55 | RSOW | 24.45 | 1.47 | 44.78 | 16.04 |
| ST59 | RSOW | 25.72 | 1.63 | 55.36 | 14.85 |

The distribution of phytoplankton showed strong differences in terms of total Chl-a and main functional groups between the two areas (Table 3). In TNB Chl-a ranged between 1.26 and 3.75 µg/L, with diatoms strongly dominating the community with a mean abundance of 87 %. In the RSOW values of Chl-a ranged between 0.29 µg/L and 2.39 µg/L, with diatoms accounting for 62 % and haptophytes for 32 % of the total biomass (Table 4). As concerns Fv/Fm in TNB values ranged between 0.2 (stations ST18 and ST23) and 0.46 (coastal station ST12), with a mean of 0.31±0.1. In RSOW, values ranged between 0.23 (station ST43) and 0.45 (station ST48), with a mean of 0.45±0.07. The analyses of phytoplankton abundance in terms of cells per liter revealed that in TNB the most abundant diatoms species were *Pseudo-nitzschia* spp. (3.1×10^5^ −1.6×10^6^ cells/L), *Fragilariopsis* spp. (5.8×10^5^ - 5.5×10^6^ cells/L), and *Chaetocheros* spp. (1.3×10^4^ - 2.8×10^5^ cells/L). Other diatoms were on average 7.4(±5)×10^5^ cells/L. *Phaeocystis antarctica* ranged between 1.6×10^5^ and 2.9×10^6^ cells/L, while the total number of dinoflagellates ranged between 1.2×10^5^ and 2.4×10^5^. Other phytoplankton groups (*Cryptophyceae*, *Crysophyceae*, *Chlorophyceae*, *Prasinophyceae* and small flagellates) reached the maximum of 3.2×10^6^ cells/L. Choanoflagellates have been reported here within phytoplankton, with a total number of cells of 1.1×10^4^ cells/L. In the RSOW, *Phaeocystis antarctica* was the most abundant species (3.2×10^5^ - 5.5×10^6^ cells/L). Among diatoms, the most abundant species in RSOW were *Pseudo-nitzschia* spp. (1.7×10^5^ - 2×10^6^ cells/L), *Fragilariopsis* spp. (1.6×10^5^ - 2.3×10^6^ cells/L), and *Chaetocheros* spp. (7.9×10^4^ - 3.1×10^5^ cells/L). Other diatoms were on average 2.6(±2)×10^5^ cells/L., while the total number of dinoflagellates showed a mean value of 1.1(±1)×10^5^. Other groups (*Cryptophyceae*, *Crysophyceae*, *Chlorophyceae*, *Prasinophyceae* and small flagellates) showed a mean value of 9.8(±3)×10^5^ cells/L. It must be noted that during the cruise, a high concentration of Cohanoflagellates was reported in the same area by (Escalera et al., 2019), with a mean total abundance of 1.0×10^6^ cells/L and a maximum of 2.9×10^6^ cells/L, two orders of magnitude higher than in TNB.

**Table 3.** Results of the pigments and chemotaxonomic analysis of the phytoplankton community of the sample stations.

| Station | Area | Chl-a<br>micro | Chl-a<br>nano | Chl-a<br>pico | Chl-c3 | Chl-c2 | Peridin<br>in | Fucox<br>anthin | 19hf* | Diadino<br>xanthin | Alloxa<br>nthin | Diatox<br>anthin | Zeaxant<br>hin | Luteine |
| --- | --- | --- | --- | --- | --- | --- | --- | --- | --- | --- | --- | --- | --- | --- |
| ST12 | TNB | 1.29 | 0.49 | 0.73 | 0.14 | 0.25 | 0 | 0.48 | 0.37 | 0.07 | 0.01 | 0 | 0 | 0.01 |
| ST14 | TNB | NA | NA | 0.15 | 0.05 | 0.47 | 0.05 | 1.43 | 0.05 | 0.18 | 0.03 | 0.01 | 0 | 0 |
| ST15 | TNB | 1.06 | 1.22 | 0.11 | 0.03 | 0.26 | 0 | 0.85 | 0.04 | 0.15 | 0 | 0 | 0 | 0 |
| ST18 | TNB | 1.33 | 0.97 | 0.09 | 0.03 | 0.26 | 0.04 | 0.82 | 0.05 | 0.14 | 0 | 0.01 | 0.01 | 0 |
| ST19 | TNB | NA | NA | NA | 0.07 | 0.31 | 0 | 0.75 | 0.11 | 0.1 | 0 | 0.01 | 0 | 0 |
| ST21 | TNB | 0.08 | 1.04 | 0.18 | 0.02 | 0.16 | 0 | 0.48 | 0.09 | 0.09 | 0 | 0 | 0 | 0 |
| ST23 | TNB | 1.18 | 0.69 | 0.15 | 0.03 | 0.24 | 0 | 0.76 | 0.05 | 0.15 | 0.01 | 0.01 | 0.01 | 0.01 |
| ST39 | TNB | 0.79 | 0.4 | 0.08 | 0.04 | 0.15 | 0 | 0.44 | 0.1 | 0.06 | 0.01 | 0.01 | 0.01 | 0 |
| ST43 | RSOW | NA | NA | 0.08 | 0.03 | 0.36 | 0.09 | 1.05 | 0.05 | 0.17 | 0 | 0 | 0 | 0 |
| ST44 | RSOW | 0.6 | 0.24 | 0.25 | 0.06 | 0.17 | 0 | 0.27 | 0.01 | 0.04 | 0 | 0 | 0 | 0.01 |
| ST45 | RSOW | 0.13 | 0.49 | 0.12 | 0.05 | 0.12 | 0.02 | 0.15 | 0.01 | 0.05 | 0 | 0 | 0 | 0.01 |
| ST46 | RSOW | 0.38 | 1.05 | 0.2 | 0.18 | 0.32 | 0 | 0.22 | 0.05 | 0.09 | 0 | 0.01 | 0 | 0.04 |
| ST47 | RSOW | 0.41 | 0.55 | 0.26 | 0.12 | 0.02 | 0 | 0.15 | 0.03 | 0.01 | 0 | 0 | 0.01 | 0 |
| ST48 | RSOW | NA | NA | 0.11 | 0.16 | 0.24 | 0 | 0.15 | 0.04 | 0.04 | 0 | 0 | 0 | 0.02 |
| ST49 | RSOW | 1.33 | 0.86 | 0.19 | 0.12 | 0.31 | 0 | 0.49 | 0.01 | 0.07 | 0 | 0.02 | 0 | 0.01 |
| ST51 | RSOW | 0.07 | 0.13 | 0.09 | 0.08 | 0.21 | 0 | 0.34 | 0.01 | 0.05 | 0 | 0.01 | 0 | 0.01 |
| ST53 | RSOW | 0.89 | 0.07 | 0.18 | 0.1 | 0.32 | 0.04 | 0.53 | 0.27 | 0.07 | 0 | 0.01 | 0 | 0.01 |
| ST54 | RSOW | 1.29 | 0.23 | 0.17 | 0.19 | 0.45 | 0 | 0.8 | 0.4 | 0.14 | 0 | 0.01 | 0 | 0.02 |
| ST55 | RSOW | 1.87 | 0.18 | 0.17 | 0.17 | 0.39 | 0 | 0.53 | 0.59 | 0.13 | 0 | 0.01 | 0 | 0.02 |
| ST59 | RSOW | 1.41 | 0.32 | 0.23 | 0.1 | 0.27 | 0 | 0.45 | 0.02 | 0.08 | 0 | 0.01 | 0 | 0.02 |

\*19hf - 19'hexanoyl oxy fucoxanthin

**Table 4.** Photosynthetic efficiency and phytoplankton community structure in the sampled stations.

| Station | Sampling area | Fv/Fm | Chlorophyta (%) | Cryptophyta (%) | Cyanophyta (%) | Diatom (%) | Dinophyta (%) | Haptophyta (%) | Prasphyta (%) |
| --- | --- | --- | --- | --- | --- | --- | --- | --- | --- |
| ST12 | TNB | 0.46 | 3.96 | 1.52 | 0 | 41.95 | 12.22 | 40.35 | 0 |
| ST14 | TNB | 0.38 | 0.36 | 9.47 | 0.13 | 179.32 | 0.72 | 0 | 0.26 |
| ST15 | TNB | 0.3 | 0.33 | 0.27 | 0.12 | 115.92 | 0.72 | 0 | 0.23 |
| ST18 | TNB | 0.2 | 0.1 | 0.08 | 1.61 | 92.32 | 0 | 0 | 0.07 |
| ST19 | TNB | 0.32 | 0.35 | 0.29 | 0.13 | 104.74 | 0 | 16.81 | 0.25 |
| ST21 | TNB | 0.24 | 0.2 | 0.17 | 0.07 | 65.79 | 0 | 0 | 0.15 |
| ST23 | TNB | 0.2 | 0.21 | 2.53 | 0.08 | 97.99 | 0.25 | 0 | 0.15 |
| ST39 | TNB | 0.41 | 0.12 | 1.71 | 1.1 | 54.92 | 0 | 9.97 | 0.09 |
| ST43 | RSOW | 0.23 | 0.16 | 0.13 | 0.06 | 120.87 | 0.01 | 0 | 0.11 |
| ST44 | RSOW | NA | 4.19 | 0.1 | 0 | 44.3 | 0 | 16.93 | 0 |
| ST45 | RSOW | 0.33 | 2.83 | 0.03 | 0 | 16.8 | 4.64 | 19.84 | 0 |
| ST46 | RSOW | 0.3 | 12.28 | 0 | 0 | 18.47 | 0 | 80.73 | 0 |
| ST47 | RSOW | 0.28 | NA | NA | NA | NA | NA | NA | NA |
| ST48 | RSOW | 0.45 | 8.54 | 0 | 0 | 15.38 | 0 | 54.76 | 0 |
| ST49 | RSOW | 0.42 | 4.36 | 0 | 0 | 49.24 | 0 | 46.15 | 0 |
| ST51 | RSOW | 0.3 | 2.83 | 0.09 | 0 | 42.7 | 0 | 33.88 | 0 |
| ST53 | RSOW | 0.26 | 2.84 | 0.04 | 0 | 58.23 | 0 | 26.14 | 0 |
| ST54 | RSOW | 0.36 | 5.4 | 0 | 0 | 79.83 | 0 | 56.82 | 0 |
| ST55 | RSOW | 0.3 | 7.15 | 0.05 | 0 | 57.25 | 0 | 44.01 | 0 |
| ST59 | RSOW | 0.27 | 5.48 | 0.13 | 0 | 57.53 | 0 | 41.01 | 0 |

### Diversity of the bacterial community

Bacterial diversity was investigated using the 16S rRNA gene sequence. After quality check and data filtering, a total of 535,009 reads were obtained and used to identify 703 unique Amplicon Sequence Variants (ASVs). Simpson and Shannon diversity index showed higher diversity in the stations of RSOW than TNB (Figure 2), albeit the differences were not statistically significative (Kruskal-Wallis test). Despite these differences, the bacterial community structure at the phylum level was similar among the stations of the two areas. This apparent similarity is still visible all the way to the family level (Figure 3). Sequences belonging to the *Bacteroidetes* and *Proteobacteria* represented the most abundant phyla in all the samples, with on average 50.1 % of the reads assigned to the *Bacteroidetes* and 48.4 % to the *Proteobacteria* respectively. In addition, sequences classified as belonging to the phyla *Cyanobacteria, Firmicutes, Actinobacteria* and *Verrucomicrobia* were detected in almost all samples.

**Figure 2.**
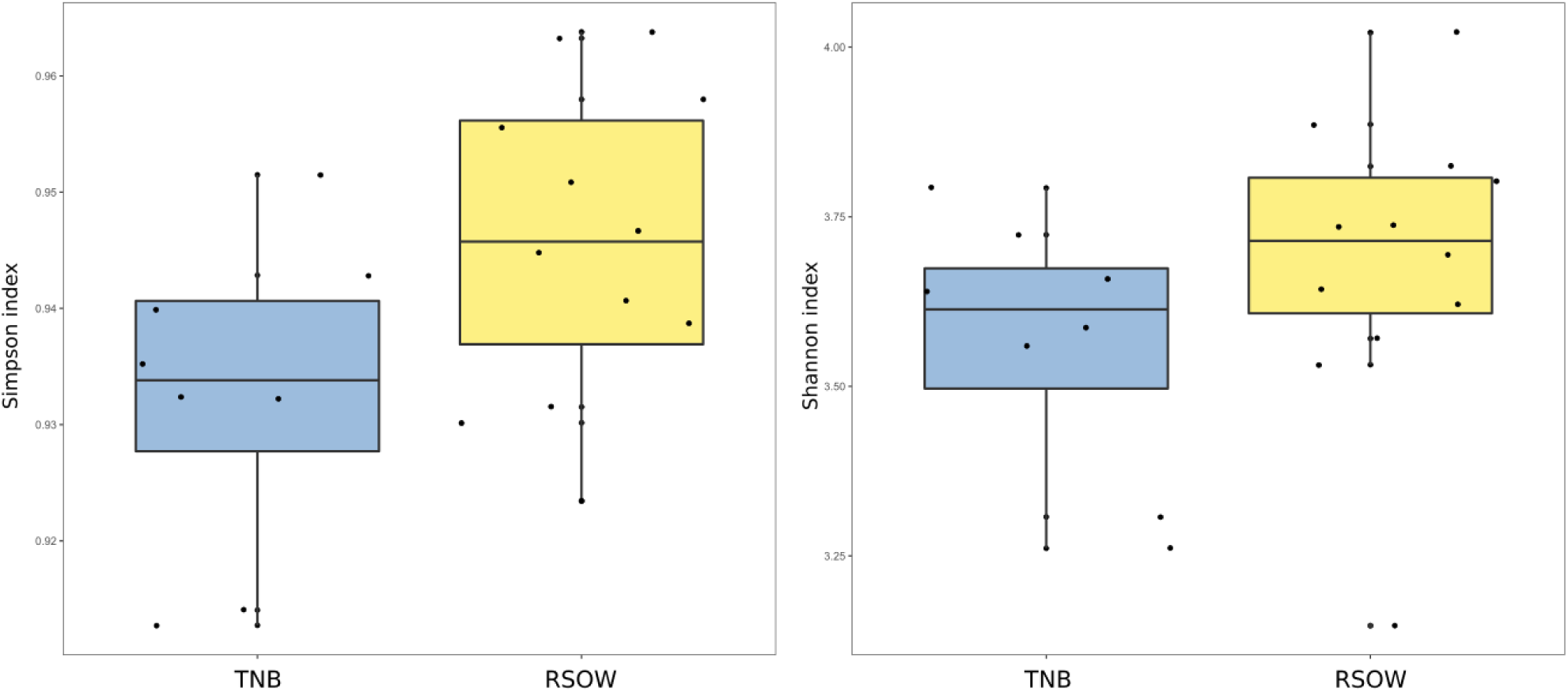
Alpha diversity metrics across the two sampled areas. a) Simpson diversity index and b) Shannon diversity index.

**Figure 3.**
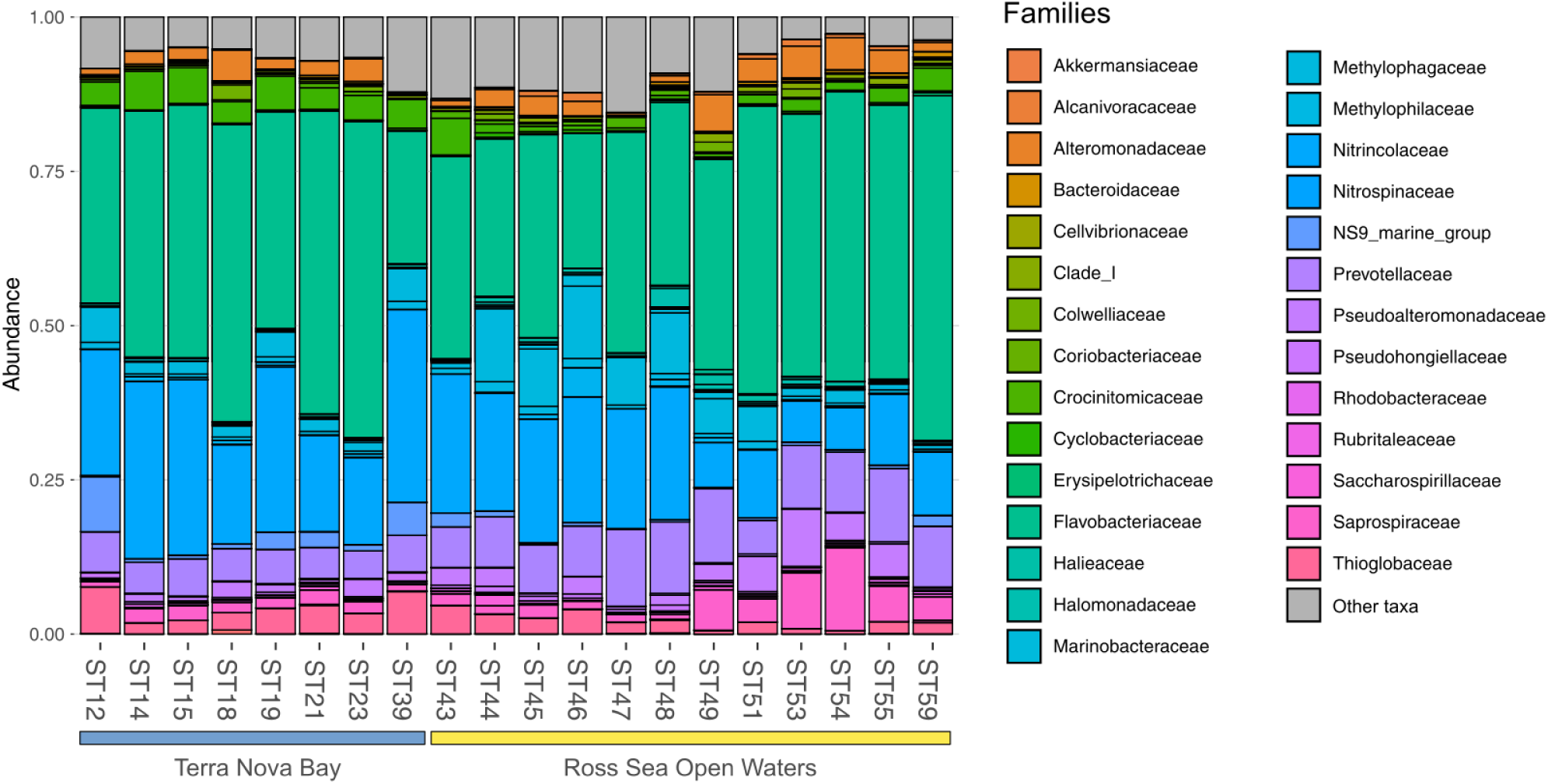
Family level distribution of the 16S rRNA diversity at the deep-chlorophyll maximum of the sampled stations. Only the most abundant families are reported, while the rare taxa are grouped together in the *Other taxa* category. Stations are grouped with an horizontal bar in two distinct clusters representing the Terra Nova Bay stations and the Ross Sea Open Water stations. Relative abundance reported as 1 =100% of the total reads.

The phylum *Bacteroidetes* was dominated by the class of *Bacteroidia,* with the *Flavobacteriales* representing the most abundant order. Among *Flavobacteriales*, several ASVs were assigned to the genera *Polaribacter*, *Aurantivirga*, and *Brumimicrobium*, representing the top genera in the studied area (Figure 4A). Within the *Proteobacteria*, most sequences were affiliated to the class *Gammaproteobacteria* (47.41 %), abundant both in the coastal (TNB) and in the offshore (RSOW) stations, followed by a lower percentage of ASVs assigned to the class Alphaproteobacteria (about 1% of total ASVs). *Gammaproteobacteria* were mainly represented by the orders *Oceanospirillales,* more abundant in the stations of the TNB area than in RSOW area (23.59 % of ASVs found in TNB versus 16.74 % found in RSOW), followed by *Cellvibrionales* and *Alteromonadales,* which instead were more abundant in the offshore area (Figure 4B). Within the *Oceanospirillales,* several species were affiliated with the genera *Alcanivorax*, *Oleispira*, *Halomonas* and *Profundimonas*, all known hydrocarbon degraders, distributed almost all the sampled stations, while among the *Cellvibrionales* the main representatives were members of the clades SAR92 and OM60(NOR5). In addition, among *Alteromonadales* we found mainly sequences classified as members of the *Pseudoalteromonas*, *Marinobacter* and *Colweilla* genera. *Alphaproteobacteria* were mainly represented by the order SAR11_Clade and *Rhodobacterales*, uniformly distributed in both the studied areas representing on average 0.55 % and 0.33 % of the total reads, respectively (Figure 4C)

**Figure 4.**
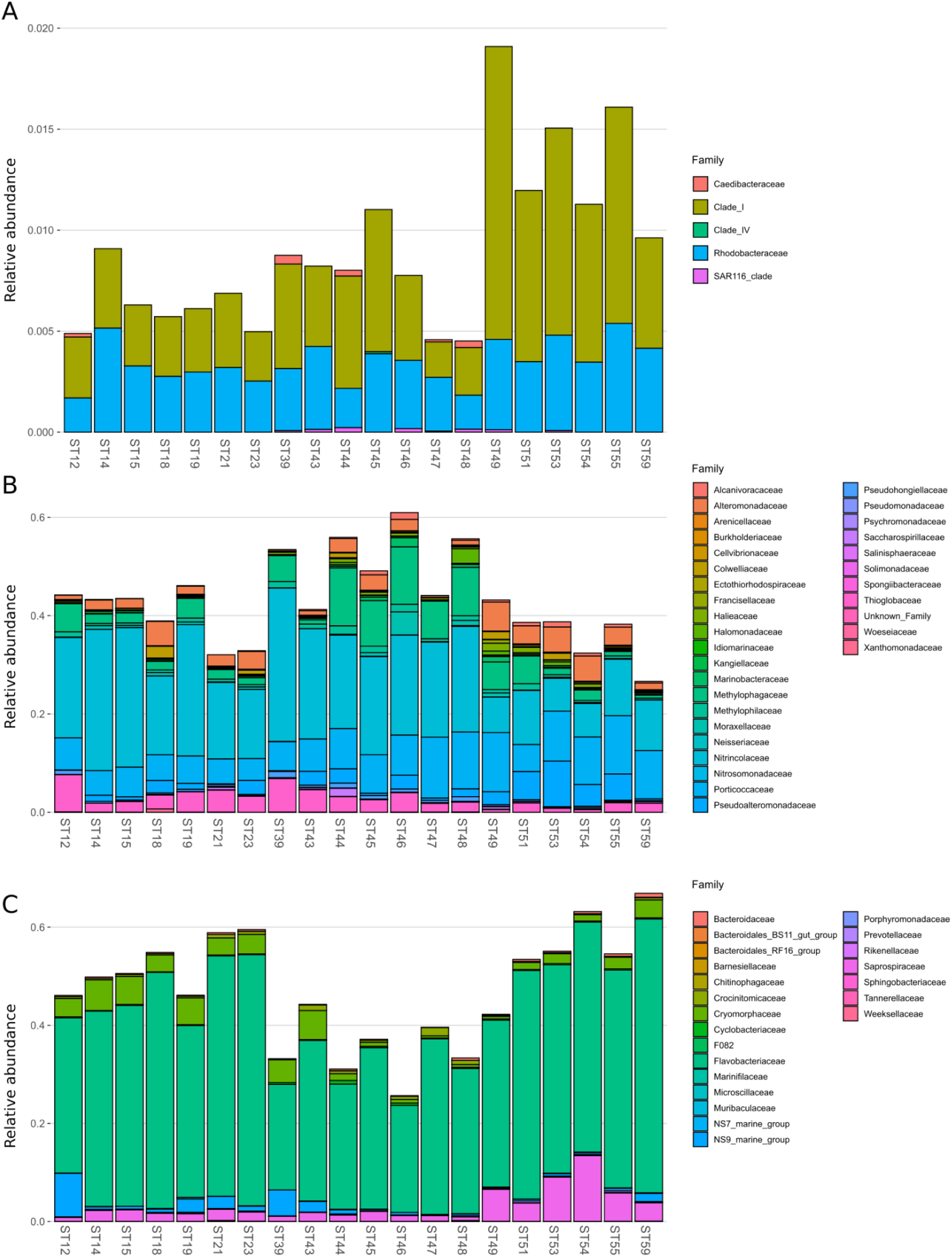
Family level distribution of the 16S rRNA diversity within the class Alphaproteobacteria (A), Gammaproteobacteria (B) and Bacteroidia (C) for the sample stations.

While the abundance of possible hydrocarbonoclastic bacteria was relatively low, representing at most 3 % of the total community (Figure 5), they were distributed differentially among the sampled areas. As shown in Figure 5, members of the genera *Alcanivorax*, *Marinobacter* and *Oleispira* were present in all the sampled stations, with a higher abundance in the offshore stations of RSOW with, on average, 0.55 %, 0.46 % and 0.29 % of the total reads respectively. Albeit with a lower abundance, we found also reads classified as *Methylophaga* and *Methylobacillus* in several stations of the RSOW area, while completely absent in the TNB stations.

**Figure 5.**
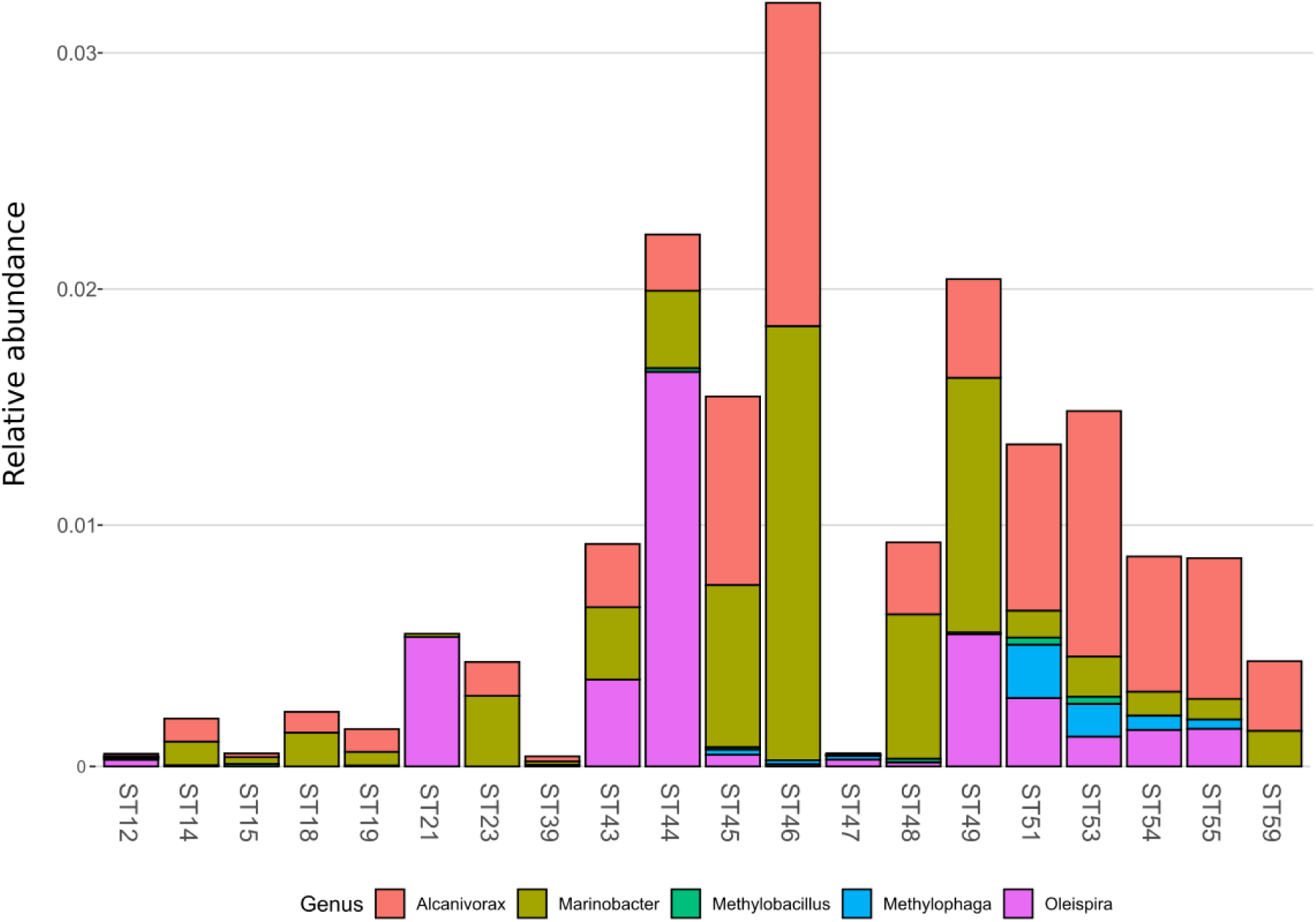
Genus level distribution 16S rRNA diversity of the putative hydrocarbon degraders detected in the sampled stations.

To explore the bacterial community composition between the stations and to identify the environmental drivers responsible for its structuring, we investigated the beta diversity distribution using both abundance weighted and unweighted dissimilarity indices. The principal coordinate analysis (PCoA) analysis using either the abundance weighted and unweighted Unifrac distance index shows a clear separation between the two studied areas in both plots (Figure 6A and B). Abundance weighted estimates of the beta diversity shows a total of 72.5 % of the variance visualized on the plot (Figure 6A), with the two study areas clearly separating along the PCoA axis 2 (explaining 21 % of the variance). The only exception to this separation is represented by station ST43, which represents the first station of the RSOW area but is geographically located near the TNB stations (Figure 1). Unweighted diversity measures show a less pronounced separation among the two sampled areas, with a lower percentage of the variance accounted for by PCoA axis 1 and 2 (cumulatively 29 % of the variance) (Figure 6B). Direct comparison of the two plots revealed that the stations had on average similar species, and that differences were due variations in the abundant species.

**Figure 6.**
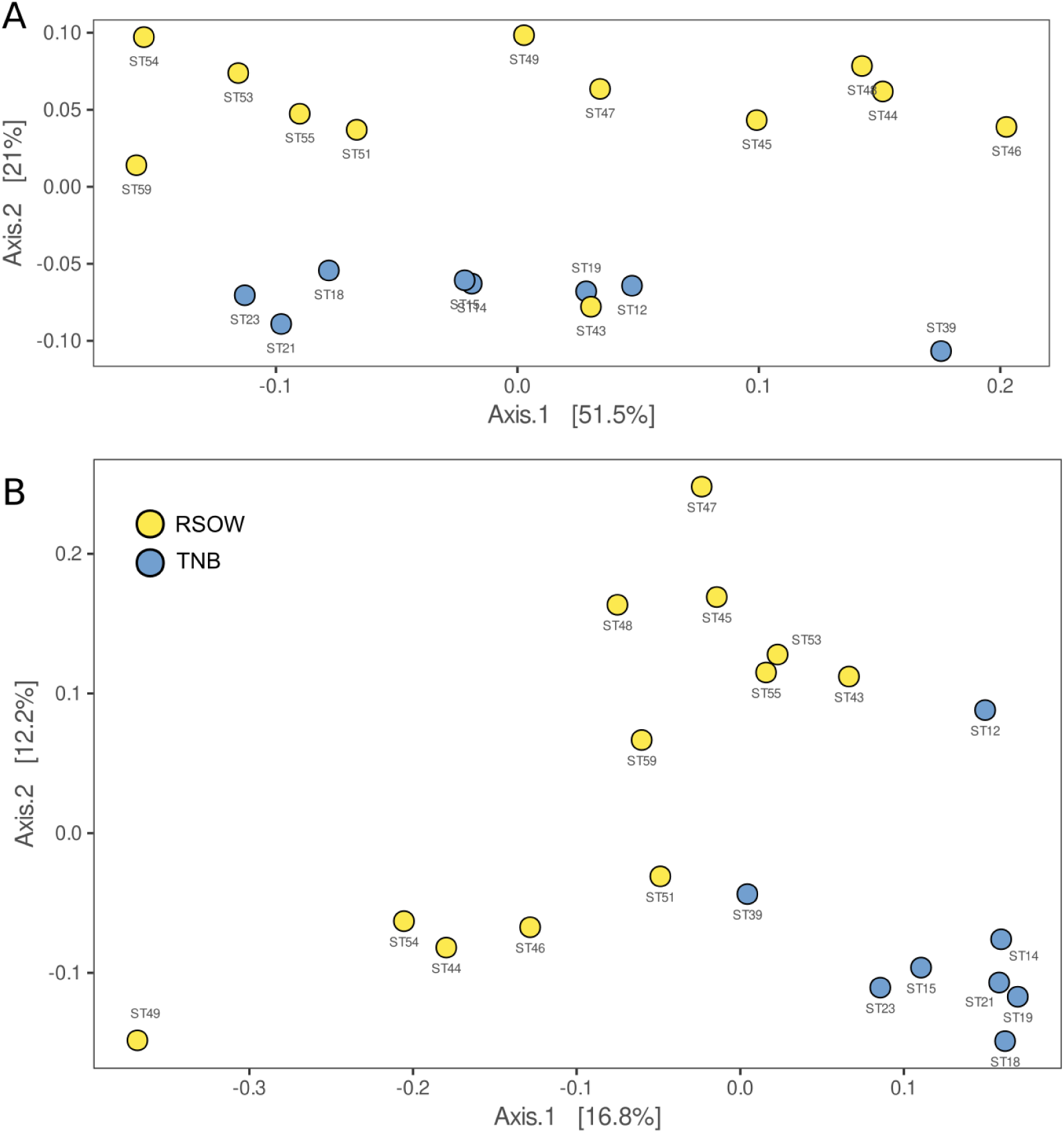
Principal coordinate analysis (PCoA) plot of the 16S rRNA gene microbial diversity based on Unifrac dissimilarity measure in the sampled stations: A) abundance weighted; and B) unweighted PCoA plots. The amount of variance explained by each axis is reported within square brackets.

To investigate the role of measured environmental parameters and the phytoplankton community composition in driving the bacterial diversity in the two studied areas, we performed non-metric multidimensional scaling (nMDS) ordination based on the abundance weighted Jaccard dissimilarity measure followed by the environmental and community composition vector fitting (Figure 7). Similarly to the results obtained with the abundance weighted PCoA analysis, the nMDS showed a clear separation among the two areas with the exception of station ST43. Linear vector fitting against the nMDS ordination revealed that the main factors explaining the bacterial diversity were the geographic location (represented by the longitude of the sampled station) and the haptophytes relative abundance, with correlation coefficients of R^2^=0.86 and R^2^=0.72 against nMDS1, respectively (Figure 7 and Table 5). The nitrogen to phosphorus ratio (N/P ratio), was instead strongly correlated with nMSD2 (R^2^=0.73), together with salinity and the maximum photosynthetic quantum yield (Fv/Fm), showing a correlation coefficient of R^2^=0.69 and R^2^=0.46, respectively.

**Figure 7.**
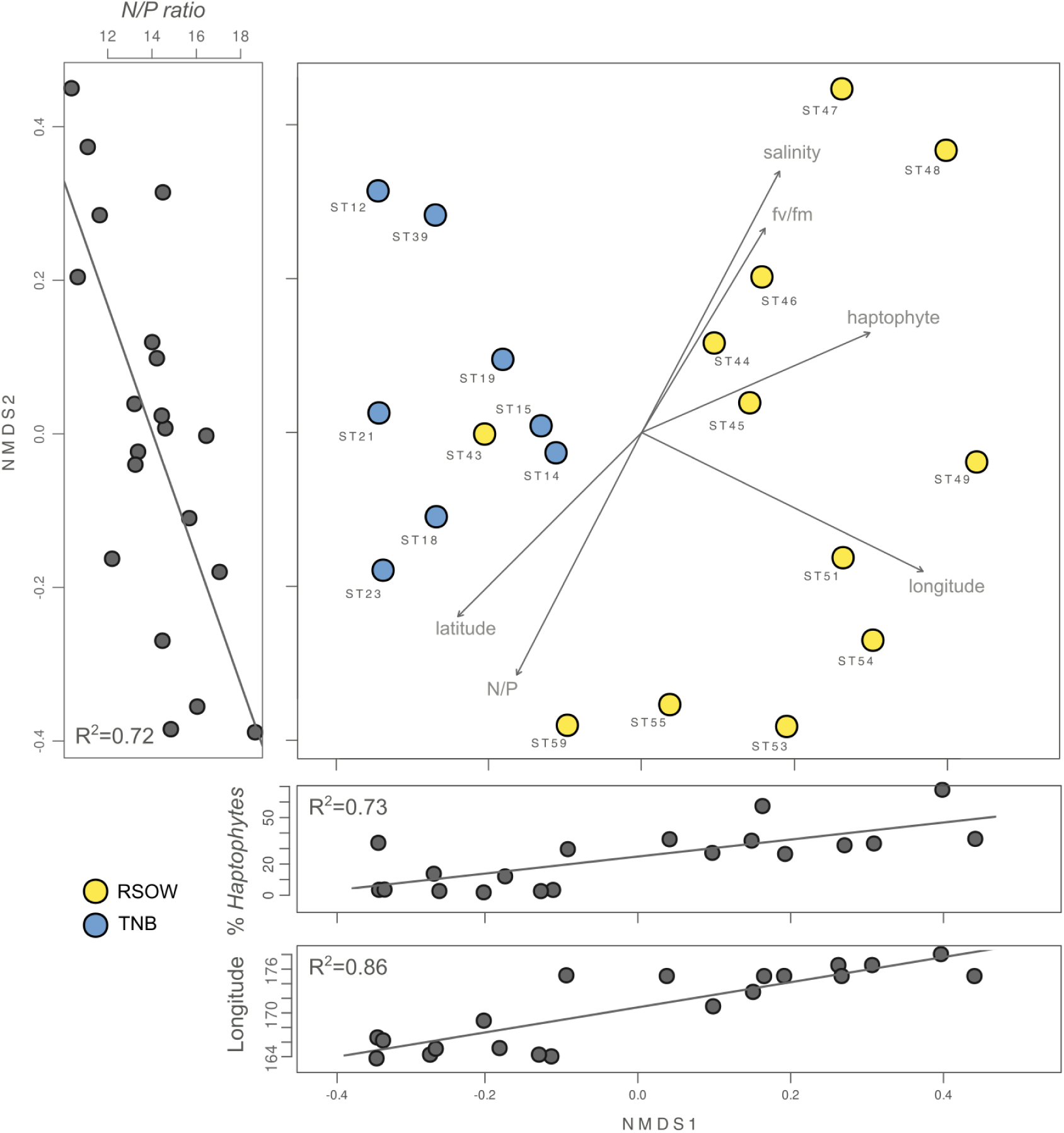
Non-metric multidimensional scaling (nMDS, stress 0.02) plot of the 16S rRNA gene amplicon microbial diversity based on Jaccard dissimilarity measure overlaid with environmental vector fitting. The lateral panels show the Pearson moment correlation (R^2^) between the respective nMDS axis and selected environmental and phytoplankton variables.

**Table 5.**
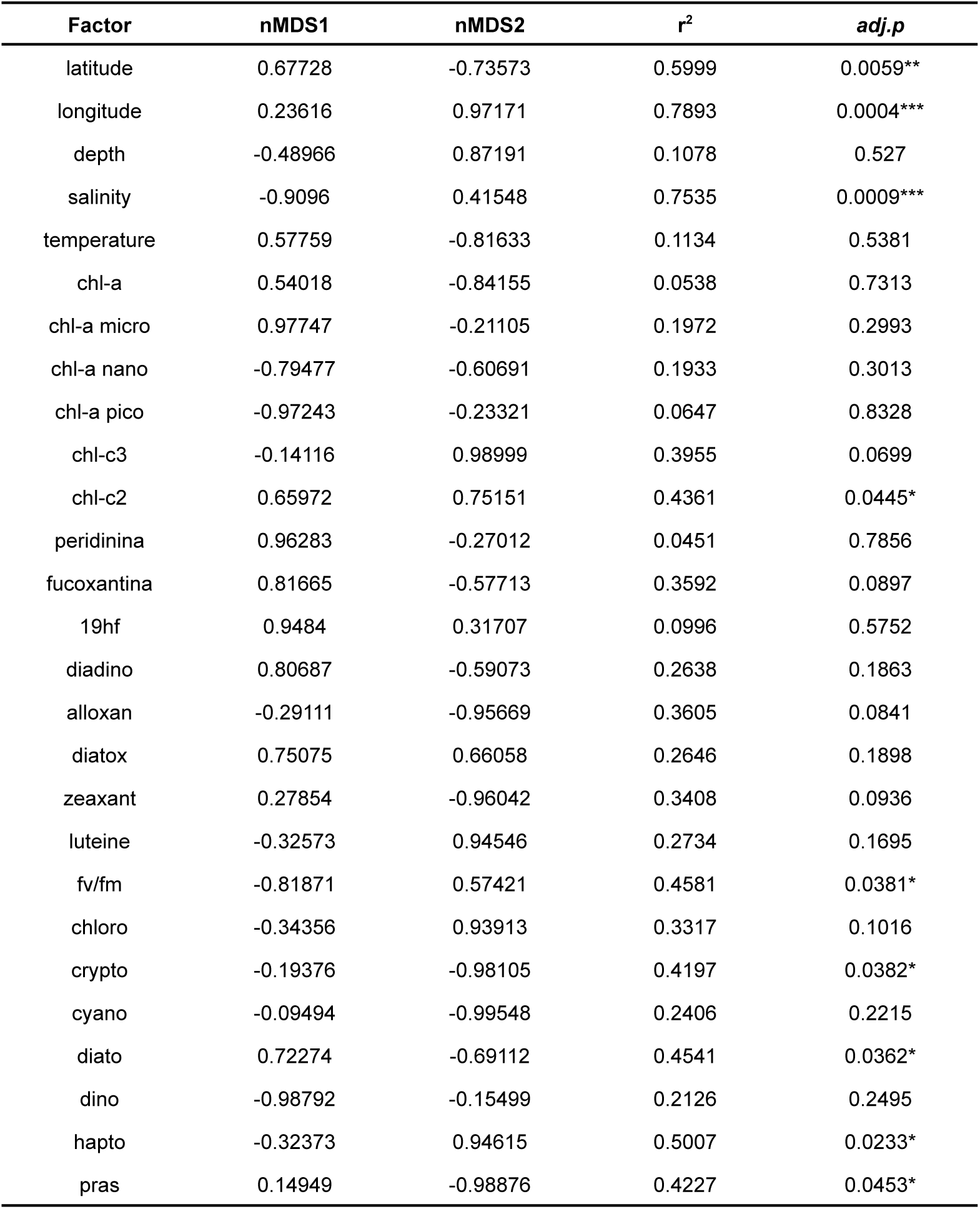
Results of the linear vector fitting of the environmental and phytoplankton community predictors on the bacterial diversity nMDS

Several other variables were identified by the linear vector fitting as significatively correlated with nMDS axis 1 and 2. This was likely due to the high degree of collinearity present among the environmental parameters and among the phytoplankton composition descriptors (Figure 8). Collinearity was investigated using Pearson moment correlation among the predictors used for the linear vector fitting, together with the results of the vector fitting for both nMDS axis (Figure 8 and Table 5).

**Figure 8.**
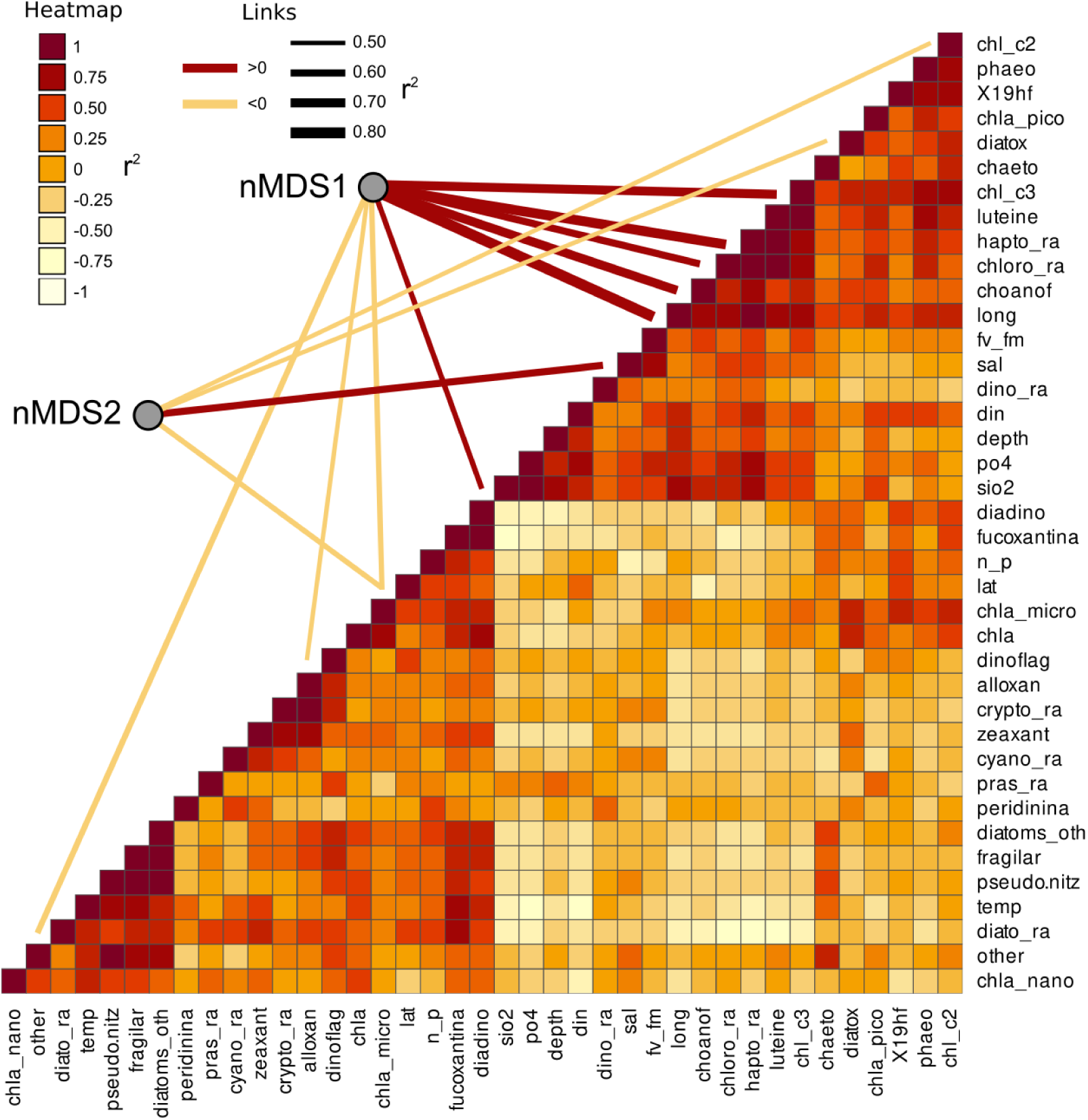
Collinearity of the environmental and phytoplankton variables used as predictors of bacterial beta diversity. The nMDS axes are connected with significant predictors through a line showing the correlation direction (color) and intensity (line thickness). Collinearity among the predictors was calculated as Pearson moment correlation and plotted as a heatmap.

### Differential distribution of key bacterial genera in the Ross Sea surface waters

The overall distribution of the bacterial community in the multivariate ordination indicates a strong role of the two areas in controlling the community assemblages, and that the separation between the two sampled areas is driven by abundant species. We determined unique and shared ASVs between TNB and RSOW area (Figure 9A), revealing that the shared community is composed of 365 ASVs, while 132 and 206 unique ASVs are found in TBN and RSOW samples respectively. We identified the top bacterial genera shared among the two studied areas (Figure 9B), and identified those showing a differential distribution. The results show several well known members of the Antarctic bacterioplankton community as the most abundant genera, some of which are differentially distributed among the two areas. Sequences belonging to the *Polaribacter* genus were highly abundant in the dataset, representing on average 47.1 % of the total reads. Among them, 41.1 % were related to *Polaribacter irgensii* (Table 6), a psychrophilic heterotrophic gas vacuolate bacterium of the *Bacteroidetes* phylum, previously isolated from Arctic and Antarctic sea ice (Gosink et al., 1998). As shown in Figure 9B, reads classified as *Polaribacter irgensii* are differentially distributed with a higher percentage of read found in the coastal area of TNB with respect to the offshore (24.3 % in TNB vs 16.8 % in RSOW, adj.p<0.01). The remaining reads are classified as members of *Polaribacter* sp. IC063 are instead uniformly distributed among the sampled areas. Members of the SUP05_cluster, a bacterial taxon comprising chemolithoautotrophs generally detected in fluids and hydrothermal plumes (Sunamura et al., 2004; Sunamura and Yanagawa, 2015) or in low oxygen areas of the water column (Walsh et al., 2009), follow a trend similar to *Polaribacter irgensii*, with the percentage of ASVs assigned to this taxon higher in the coastal area of TNB with respect to the offshore RSOW stations (4.1 % vs 2.1 %, respectively). Although with a lower abundance (less than 1 % of the total reads), we found that the ASVs associated to the genera *Profundimonas* and *Brumicrobium*, cold-adapted and facultatively anaerobic heterotrophic bacteria generally found in marine environments (Bowman et al., 2003; Margesin, 2017), are also more prevalent in the TNB samples.

**Figure 9.**
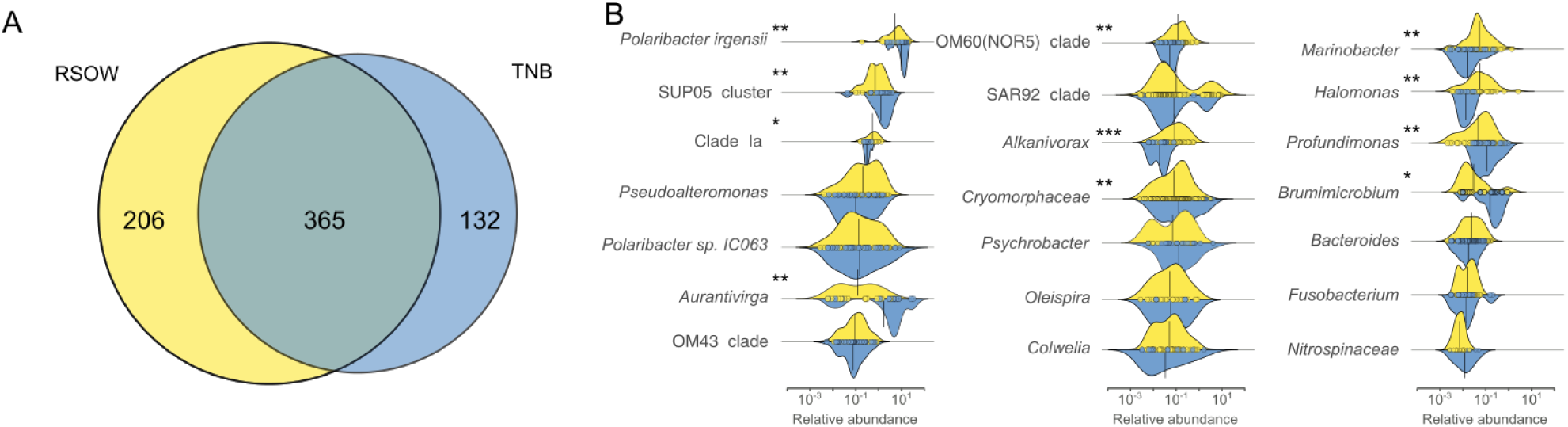
Shared and unique ASVs among the two sampled areas (A) and density function of the abundance of the top Genera shared among the sampled areas (B). Differentially abundant genera are marked with an * according to the result of the Kruskal-Wallis test (* adj.p<0.05; ** adj.p<0.01; *** adj.p<0.001).

**Table 6.**
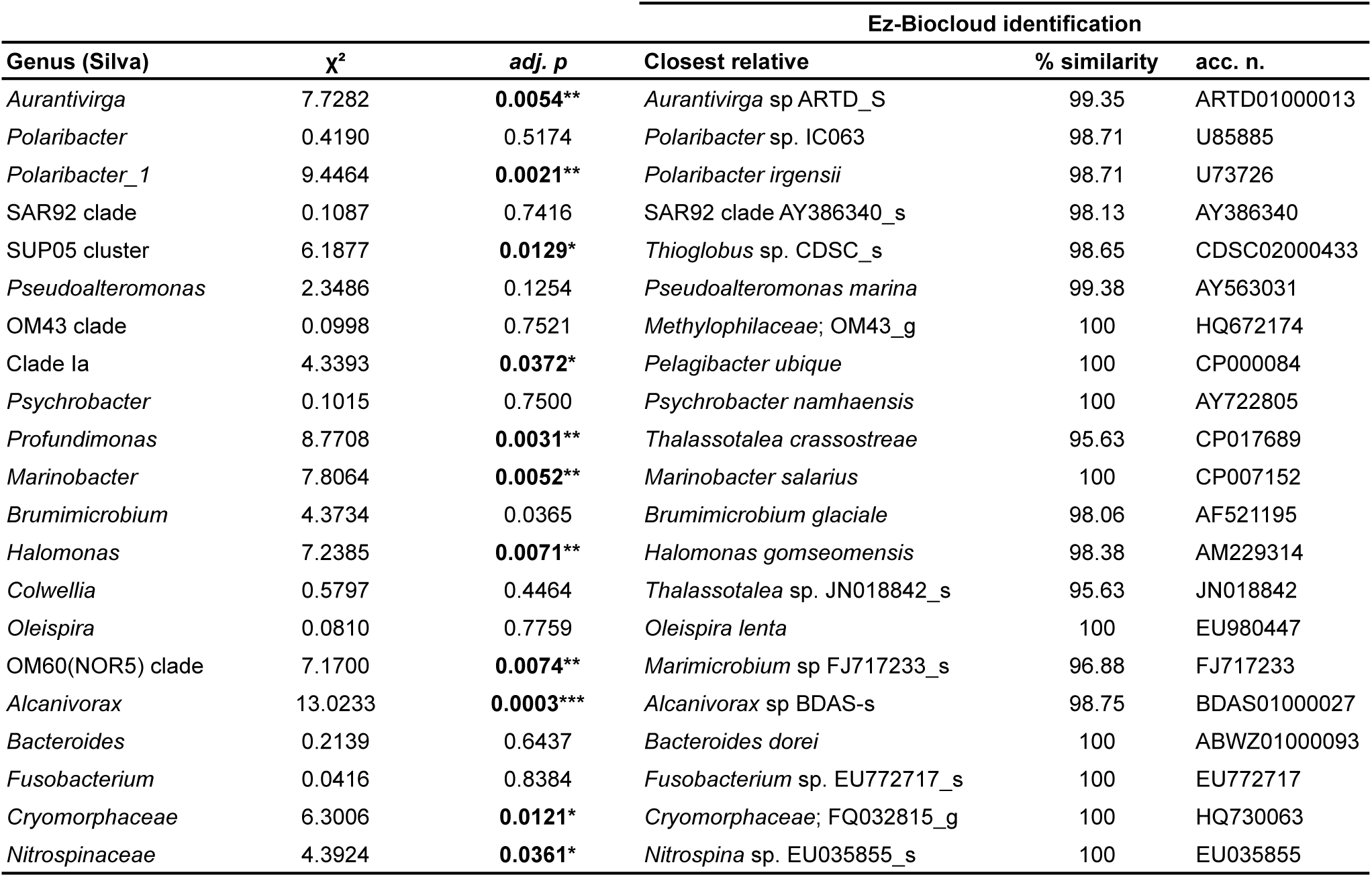
Most abundant shared genera among the two sampled areas. The table reports the results of the statistical test for differential distribution and the information relative to the closest cultured relative identified using the Ez-Biocloud identification software.

An opposite trend was followed by the reads classified as members of the genera *Aurantivirga*, and, with a markedly lower abundance, by the reads affiliated to Clade Ia, OM60 (NOR5) clade, *Alcanivorax*, *Marinobacter* and *Halomonas* (Figure 9B). *Aurantivirga* is a member of *Flavobacteriaceae* whose type strain, *Aurantivirga profunda*, has been isolated in the deep-sea waters of the Pacific Ocean (Song et al., 2015). Our data show that the ASVs belonging to *Aurantivirga* are abundant in the RSOW area (on average 4.7 %) and become substantially lower in the coastal area of TNB (0.7 %, adj.p<0.01). Our results show that the ASVs related to Clade Ia are significantly more abundant in the RSOW area (0.7 %) with respect to the TNB area (0.3 %). A similar distribution was found for members of the OM60 (NOR5) clade with a higher abundance in the offshore stations with respect to the coastal stations (0.6 % in RSOW versus 0.2 % in TNB). The genera containing obligate or facultative hydrocarbon oxidizers *Alcanivorax*, *Oleispira* and *Marinobacter* were also differentially distributed among the sampled areas, showing statistically significant higher abundances in the stations of the RSOW compared to TNB (adj.p<0.001 for *Alcanivorax* and adj.p<0.01 for *Oleispira* and *Marinobacter*).

Patterns of co-occurrence among the more prevalent ASVs (here defined as occurring in at least 20 % of the sampled stations) were investigated using different cut-off levels of Spearman correlation coefficient, between 0.5 and 0.85 (Figure 10 and Table 7). The results revealed a phylogenetically heterogeneous cluster of highly co-occurring ASVs present at low threshold of correlation (from 0.5 to 0.65 rho), progressively breaking down into small (20-30 ASVs) phylogenetically heterogeneous highly correlated clusters.

**Figure 10.**
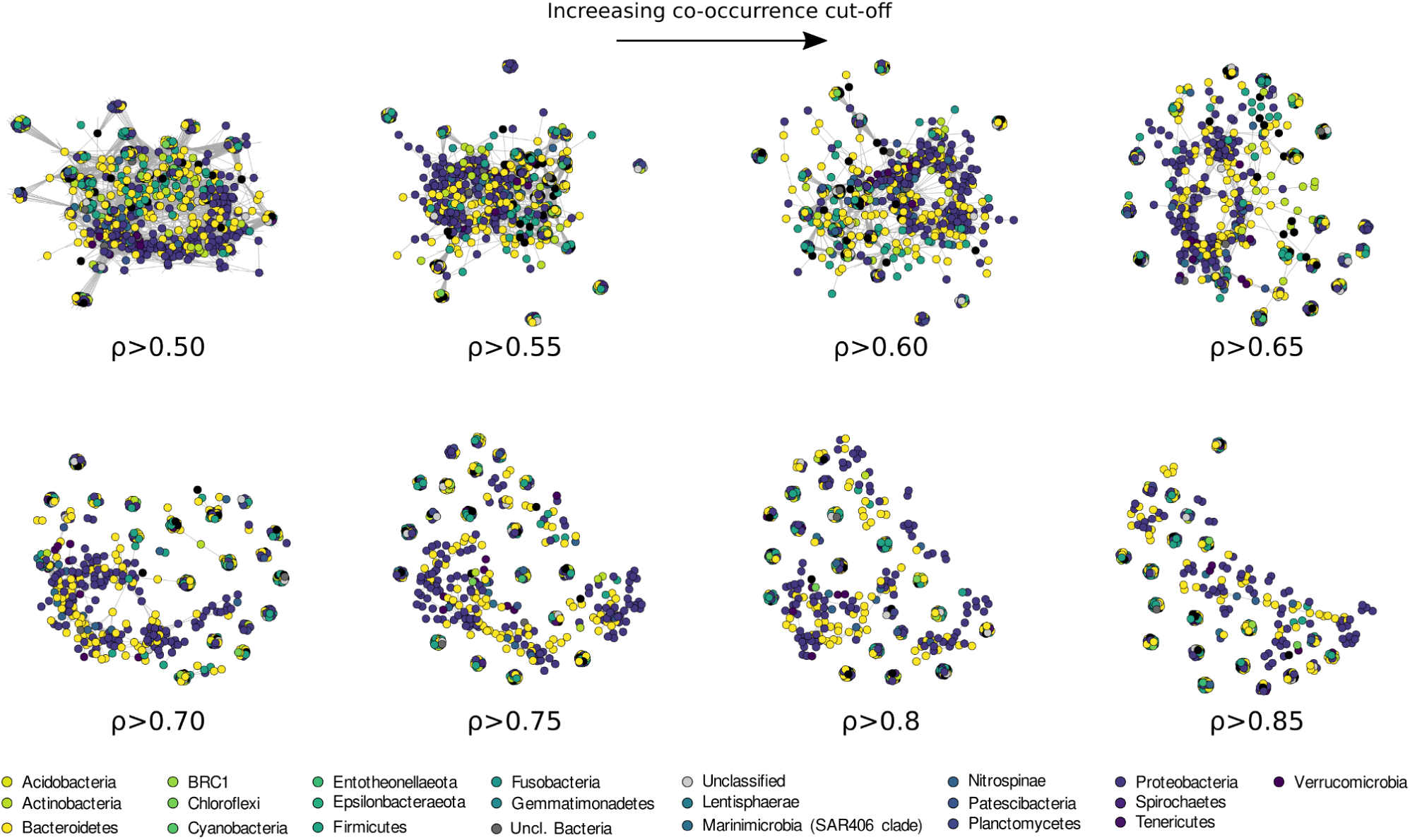
Co-occurrence network analysis drawn with increasing Spearman correlation cut off and colored according to the phyla classification of each ASVs.

**Table 7.** Network statistics for increasing correlation thresholds. SPL - Shortest path length.

| Correlation threshold ($\rho$) | Edges count | Average SPL | Assortativity | Furthest node distance | Modularity (Louvain) | Mean distance |
| --- | --- | --- | --- | --- | --- | --- |
| 0.5 | 10,124 | 1.73 | 0.48 | 4 | 0.75 | 3.23 |
| 0.55 | 7,877 | 2.44 | 0.54 | 5.51 | 0.81 | 3.74 |
| 0.6 | 6,775 | 3.2 | 0.57 | 8.67 | 0.84 | 4.43 |
| 0.65 | 4,996 | Inf | 0.74 | 12.64 | 0.91 | 7.09 |
| 0.7 | 4,342 | Inf | 0.89 | 11.72 | 0.93 | 5.69 |
| 0.75 | 3,570 | Inf | 0.97 | 11.94 | 0.94 | 4.55 |
| 0.8 | 3,270 | Inf | 0.99 | 14.13 | 0.95 | 4.63 |
| 0.85 | 3,107 | Inf | 1 | 12.29 | 0.95 | 2.01 |

## 4 Discussion

The Ross Sea is a complex mosaic of subsystems with physical, chemical, and biological features that respond to biotic and abiotic forcing at different temporal and spatial scales (Smith et al., 2007, 2014; Mangoni et al., 2017; Bolinesi et al., 2020b). Although the phytoplankton blooms dynamics in the Ross Sea have been well described, the drivers regulating the temporal and spatial distribution of the blooms are still debated (Arrigo et al., 1999; DiTullio et al., 2000; Mangoni et al., 2017, 2019; Saggiomo et al., 2021).

Among the biotic processes structuring the phytoplankton community, the interactions with the bacterioplankton has been proposed to be an important factor (Richert et al., 2019). Despite the bacterioplankton composition in the Ross Sea has been previously investigated (Lo Giudice et al., 2012; Silvi et al., 2016), the phytoplankton-bacterioplankton co-occurrence in the area has been poorly characterized. Many studies have suggested the existence of sophisticated ecological interactions between phytoplankton and bacteria in driving marine biological processes, which can range from a mutualistic exchange of biomolecules and nutrients to the competition for the same limiting inorganic nutrients (Amin et al., 2012, 2015; Seymour et al., 2017). In this study we have investigated the bacterial diversity of the Ross Sea in relation to phytoplankton community structure in 20 stations located between the coastal area of Terra Nova Bay (TNB) and the offshore waters of the central Ross Sea (RSOW). The two areas were characterized by different amounts of phytoplankton biomass and dominant functional groups, with highest levels of biomass near the coastline. *Pseudo-nitzschia* spp. and *Fragilariopsis* spp. were the most abundant species in TNB, while *Phaeocystis antarctica* dominated the stations in the RSOW. Beyond the differences in terms of species composition, the two areas differed also in terms of biomass, that was one order of magnitude higher in TNB.

The bacterioplankton community was representative of Antarctic surface waters in accordance with previous studies (Abell and Bowman, 2005; Ghiglione and Murray, 2012; Grzymski et al., 2012; Wilkins et al., 2013), with *Bacteroidetes* and *Proteobacteria* representing the most abundant phyla. Among *Bacteroidetes*, members of the *Flavobacteriaceae*, commonly found bacteria in polar environments (Abell and Bowman, 2005; Williams et al., 2012), dominated both the sampled areas. These bacteria are generally classified as the key degraders of phytoplankton derived organic matter (Teeling et al., 2012; Buchan et al., 2014). Their presence in all the sampled stations suggests that these bacteria might play an important role in the organic carbon cycle of the Ross Sea, potentially impacting organic carbon transfer to higher trophic levels (Azam and Malfatti, 2007). Within our dataset, members related to the *Polaribacter* genus were dominant in all the sampled stations, and showed a statistically higher abundance in the TNB area (Figure 9B). The closest relative to our sequences was *Polaribacter irgensii*, with an average similarity of 98.71 % (Table 6), a known psychrophilic heterotrophic marine bacterium. Previous studies have reported that *Polaribacter* species thrive during diatom-bloom (Teeling et al., 2012; Williams et al., 2012), potentially suggesting that their presence in the TNB area might be linked with the higher presence of diatom species we have identified. Similarly, a recent analysis revealed that species of the *Aurantivirga* genus are among the firsts prokaryotic taxa responding to diatom bloom in the Southern Ocean (Liu et al., 2020). Sequences related to the genus *Aurantivirga* are highly abundant in our TNB stations, suggesting once again a possible relationship with the high abundance of diatoms (Figure 9B). Among the *Proteobacteria*, sequences related to the *Gammaproteobacteria* were detected in all the sampled stations, with a differential distribution of members related to the *Alteromonadales*, *Cellvibrionale*s *and Oceanospirillales* orders among the two studied areas. Within the *Alteromonadales,* sequences related to *Pseudoalteromonas*, *Marinobacter* and *Colweilla* genera were dominant and in higher abundance in the RSOW stations (Figure 9B). These bacterial genera comprise cold-adapted marine bacteria generally detected in the Southern Ocean waters able to degrade simple sugar, amino acids, organic compounds and hydrocarbons (Médigue et al., 2005; Methé et al., 2005; Singer et al., 2011; Rosenberg, 2013). The abundance of ASVs belonging to this psychrophilic bacterial group in the offshore area could be explained by specific adaptation to the conditions found in offshore stations. Similarly, members of the *Cellvibrionales* order dominated the offshore area. These bacteria, generally affiliated to the Oligotrophic Marine Gammaproteobacteria (OMG) cluster (Cho and Giovannoni, 2004), are able to adapt to nutrient depletion and carbon limitation with the potential to harvest light for mixotrophic growth (Stingl et al., 2007; Spring and Riedel, 2013; Courties et al., 2014). Within our dataset, we also detected sequences related to the clades SAR92 and OM60 (NOR5). Previous reports indicate that members of SAR92 clade dominate during the phytoplankton bloom in the Southern Ocean (West et al., 2008) and are able to establish a close relationship with the productive *P. antarctica* during the austral summer in the Amundsen sea polynya, suggesting an important role of this bacterial taxa during bloom formation and bloom longevity (Delmont et al., 2014). Consistent with this data, our results show a dominance of the SAR92 clade in the RSOW area which is dominated by *P. antarctica*. The ability to exploit different metabolic pathways based on the conditions found in the sea water, suggest that both *Alteromonadales* and *Cellvibrionales* can play an important role in the microbial food web, contributing to the functioning of the Antarctic marine ecosystems. Taken together these observations suggest a tight coupling between the phytoplankton community structure and the dominant bacteria identified in the Ross Sea surface waters.

Among the top genera identified in our study, members belonging to the SAR11 order of the *Alphaproteobacteria* Clade Ia, were comparatively low in relative abundance. Our results show that ASVs classified in this group were more abundant in the RSOW area. Species belonging to the SAR11 order comprise aerobic and free-living oligotrophic chemoheterotrophic bacteria globally distributed in marine environments (Morris et al., 2002). They are believed to contribute significantly to the carbon, nitrogen and sulfur cycling in the Ocean (Malmstrom et al., 2004), and have been previously reported worldwide (Field et al., 1997; Morris et al., 2002; Brown et al., 2012). Members of Clade Ia represent a specific subgroup of the order generally reported in cold waters (Brown et al., 2012; Delmont et al., 2019). The closest relative to our sequences was *Pelagibacter ubique*, the most common heterotrophic bacteria found in the ocean (Giovannoni, 1990; Giovannoni et al., 2005), with an average similarity of 100 % (Table 6). Studies based on genome sequences analysis and *in situ* hybridization, revealed that *P*. *ubique* is a bacterium with a small genome size and a high metabolic activity, able to assimilate either dissolved free amino acids and dimethylsulfoniopropionate (DMSP) (Malmstrom et al., 2004; Giovannoni and Stingl, 2005). DMSP is an organosulfur compound produced by several phytoplankton cells, which can perform a double function in polar microalgae, as osmolyte or cryoprotective agent (Kirst et al., 1991; Karsten et al., 1996). Several reports indicate that *P*. *antarctica* is one of the leading producers of DMSP in the Ross Sea (DiTullio and Smith, 1995; Ditullio et al., 2003).

Interestingly, our dataset reveals the presence of obligate or facultative hydrocarbon oxidizers in the Ross Sea surface waters, albeit at abundances below ∼3 %. While the presence of facultative hydrocarbon oxidizers *per se* is not indicative of hydrocarbon contamination in the environment, our data reveal that the obligate hydrocarbonoclastic genera *Alcanivorax* and *Oleispira* (Yakimov et al., 1998, 2003) were present in all the sampled stations, and markedly more abundant in the RSOW stations. *Alcanivorax* and *Oleispira* are two bacterial genera found globally in extremely low abundances (Cafaro et al., 2013), but can become transiently dominant, with relative abundances up to 70-90% of prokaryotic cells, in the presence of hydrocarbon spills (Kasai et al., 2002a, 2002b). Their presence in our dataset is interesting and can be explained in several different ways. While their abundance is significantly lower than reported after oil spillage events (Kasai et al., 2002b; Harayama et al., 2004), it is possible that hydrocarbon and exhaust oils released by the large number of ships transiting the Ross Sea every summer are responsible for keeping them above the normal background levels. Antarctic tourism has been steadily increasing over the years, with over half a million tourist landings reported for the 2017-2018 season (Palmowski, 2020), and fishing activities in proximity of the productive Antarctic waters have also increased (Brooks and Ainley, 2017). Alternatively, previous studies have proposed that members of the *Alcanivorax* genera are able to maintain viable status in uncontaminated marine waters degrading natural lipids of bacterial and phytoplankton origin, released in the water column due to exudation, sloppy feeding or viral lysis (McGenity et al., 2012; Lea-Smith et al., 2015; Zadjelovic et al., 2020). During our sampling, the phytoplankton community was undergoing a summer bloom, and thus all the mentioned processes, *e.g.* exudates, sloppy feeding by the zooplankton grazers and viral induced lysis, might have contributed in increasing lipid concentrations in seawater. In addition to *Alcanivorax* and *Oleispira*, the facultative hydrocarbon degraders *Marinobacter* and *Halomonas* have been also identified (Figure 5). Members of the genus *Marinobacter* are slightly or moderately halophilic, able to degrade both aliphatic and aromatic hydrocarbons. *Marinobacter* spp. capable of growing on hydrocarbons as the sole carbon source have been previously isolated from sediments (Gauthier et al., 1992). The genus *Halomonas* comprises marine halophilic and/or halotolerant bacteria, known to produce large quantity of exopolysaccharides (EPS) with rheological and active-surface properties (Calvo et al., 1998, 2002; Martínez-Checa et al., 2002; Pepi et al., 2005; Gutierrez et al., 2020). It’s possible that Halomonas-producing EPS provides a mechanism to increase the bioavailability of hydrophobic compounds (e.g. hydrocarbons and lipid aggregates) that *Alcanivorax*, *Oleispira and Marinobacter* utilize for growth. Information regarding the structure of bacterial communities can be also investigated through the use of network analysis. Most microbial network analysis uses a single correlation threshold to identify meaningful interactions from a correlation matrix (Barberán et al., 2012; Williams et al., 2014; Giovannelli et al., 2016; Connor et al., 2017). This approach, while might provide useful information regarding the biological interactions in the community, requires *a priori* justification or a sensitivity analysis to demonstrate the robustness of the conclusion with respect to the selected threshold. In addition, single threshold co-correlation analysis is considered controversial by some authors (Hirano and Takemoto, 2019; Blanchet et al., 2020) and it is believed to increase the number of detected false positives. An alternative approach to identifying biological interactions is using co-occurrence to identify ecological response to common environmental factors, an approach recently applied with success across broad ecological gradients (Delgado-Baquerizo et al., 2018; Fan et al., 2018; Fullerton et al., 2021). Microbial co-occurrence networks can be also used to investigate community assembly and dynamics. To this end, we used an increasing correlation threshold to identify the structuring of the bacterioplankton community in the surface waters of the Ross Sea, preserving the most ecological signal compared to a fix-threshold approach (Figure 10). With this approach, the degree at which the network breaks apart and changes in structure during thresholding can be used to identify processes influencing community assembly (Connor et al., 2017). As the threshold is increased, the largest network component becomes sparser as edges among nodes are removed (Figure 10). If modularity and other network statistics follow a linear trend during thresholding, this suggests the presence of a dominant core microbiome, often proposed to be critical or keystone components of the community (Faust and Raes, 2012; Mandakovic et al., 2018). Conversely, non-linear trends in network modularity suggest the absence of representative core microbiome and possibly the absence of keystone species.

Our data show that the bacterioplankton community is composed of phylogenetically heterogeneous clusters of bacteria showing progressively lower co-occurrence as the correlation threshold is raised (Figure 10). Usually, high levels of co-occurrence in microbial networks are interpreted as increasing biotic interactions among species, including competition for resources (Faust and Raes, 2012; Widder et al., 2014). The resulting sparse network we have obtained at higher level of correlation threshold (>0.60 rho, Figure 10 and 11, Table 7), suggests that stochastic processes (e.g. neutrality, dispersion, and physical stress, see Holyoak et al., 2005), rather than competition, are responsible for community assemblage in the surface waters of the Ross Sea. Additionally, our results suggest the absence of a defined core microbiome in the studied area. This might be connected to different bacterial assemblages being associated to distinct phytoplanktonic communities identified in the Terra Nova Bay and the Ross Sea Open Water areas. Despite this, the presence of small clusters (20-30 ASVs) of phylogenetically diverse but recurrent ASVs suggest a strong functional redundancy among the most representative bacterial species. Ecological network theory predicts that communities of tightly connected species should be more fragile. Our data suggests that the higher fragmentation identified might be the dynamic response of the bacterial community to phytoplankton related dynamics, which continuously rework and redistribute microbial niches as blooms succession progresses (Luria et al., 2017)

**Figure 11.**
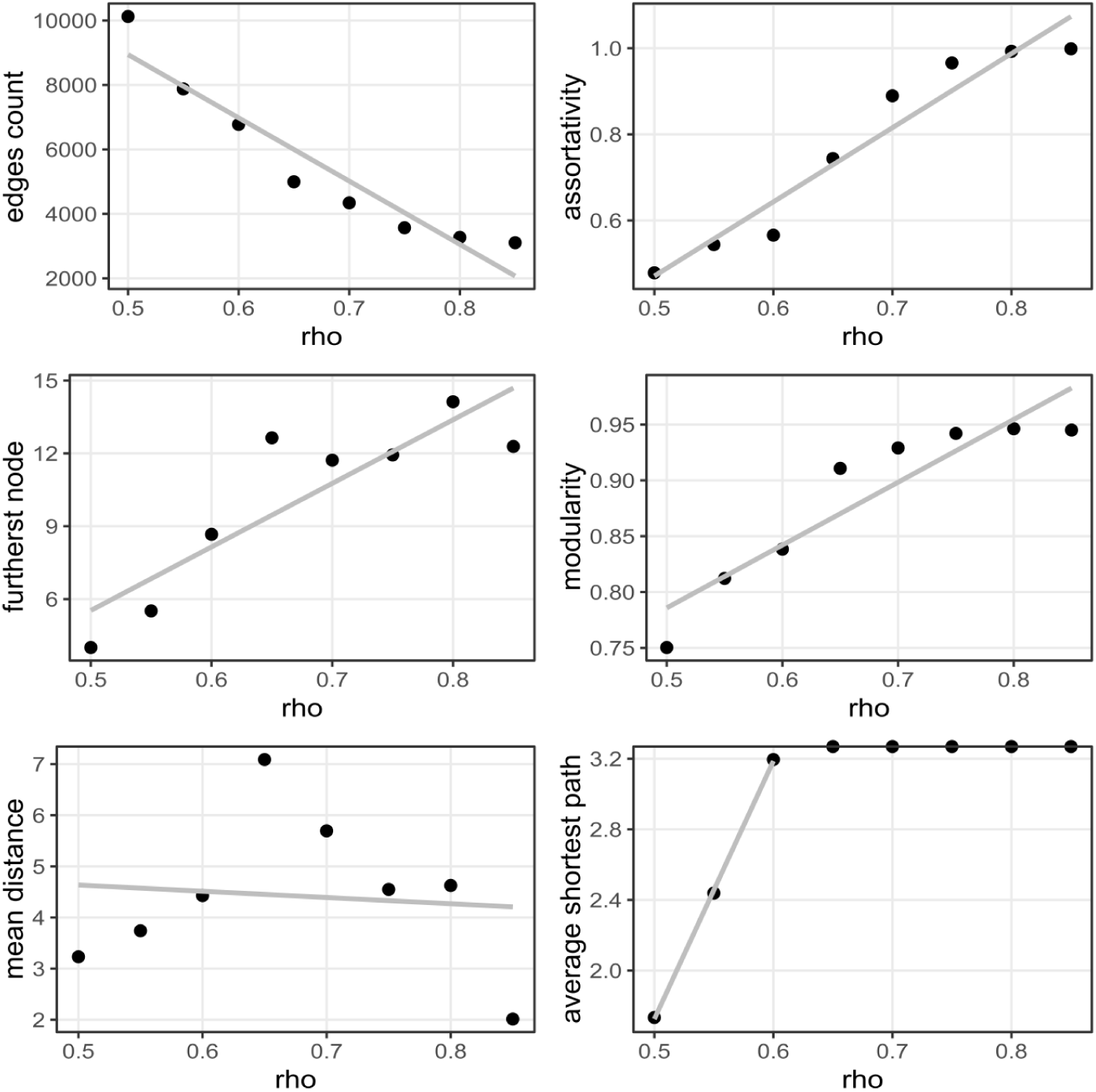
Changes in network statistics as a function of increasing correlation thresholds. A linear model was fitted to the data (grey line). A clear change in network properties is present for correlation thresholds ≥ 0.60.

### Conclusions

Microbial community distribution and diversity in marine habitats, such as polar environments, is a key factor in structuring the ecosystem functioning. Ross Sea is one of the most productive places where interaction between bacterioplankton and phytoplankton seems to strongly contribute to the ocean carbon cycle and biogeochemical processes. Our results suggest a strong connection between the phytoplankton community and the dominant bacteria identified in the Ross Sea surface waters. We found that *Bacteroidetes* and *Proteobacteria* are the most abundant phyla in all the sampled stations, with a dominance of heterotrophic bacterial species (i.e. *Polaribacter* genus) in response to diatom blooms in the TNB area and a dominance of oligotrophic and mixotrophic bacterial species (*i.e.* the OMG group and SAR11) in response to haptophytes blooms in the offshore area. Moreover, our data suggests the presence of functional redundancy within the bacterioplankton community. Our results emphasize the ability of bacteria to easily adapt to the environmental changes encountered, and suggest the existence of a specific rather than random interaction between dominant phytoplankton-bacterioplankton species that contributes to the organic matter cycling in the Southern ocean and more in general to the Antarctic food web.

## 6 Author Contributions

OM, AC and PR designed the study. MM, AC, PR, MS, FB, OM performed laboratory analyses. MM, AC, DG, MS, MB, RDM and GD performed bioinformatic and statistical analyses. All authors contributed equally to the manuscript writing and revision.

## Acknowledgments

We express our gratitude to Italian Antarctic National Program (PNRA) and the scientific personnel and crew of the research vessel Italica for logistical support.

## Data Availability Statement

The datasets generated for this study can be found on the Github repository https://github.com/giovannellilab/Cordone_et_al_Ross_Sea, permanently stored with the ZENODO under DOI https://doi.org/10.5281/zenodo.4784454. Sequencing data have been deposited in the EMBL ENA repository with accession number ERP129169. The script containing all the bioinformatic and statistical code to reproduce the results is available on the same github and DOI reported above.

